# The structural and functional modularity of ovarian follicle epithelium in the pill-millipede *Hyleoglomeris japonica* (Myriapoda: Diplopoda: Pentazonia: Glomerida) and the morphological reconsideration of somatic tissues around oocytes in Myriapoda

**DOI:** 10.1101/2023.05.17.541077

**Authors:** Yasuhiko Chikami, Kensuke Yahata

**Affiliations:** Graduate School of Life and Environmental Sciences, University of Tsukuba, Tsukuba 305-8572, Japan; Faculty of Life and Environmental Sciences, University of Tsukuba, Tsukuba 305-8572, Japan

**Keywords:** Ovarian follicle epithelium, Oogenesis, Myriapoda, Arthropoda, Modularity

## Abstract

Ovarian somatic tissues around oocytes are essential for oogenesis in many metazoans. The functional ovarian somatic tissues sometimes have structural properties specific to their roles. In contrast, there is no evidence of such structural modularity of the follicle epithelium in Myriapoda, suggesting that myriapod ovarian soma may not participate in oogenesis. The ultrastructural nature of follicle cells also supports solitary oogenesis in most myriapods investigated. In contrast, here, we report two structurally and developmentally distinct domains of the follicle epithelium in the Japanese pill millipede, *Hyleoglomeris japonica*. The follicle epithelium of *H. japonica* has a thick cell mass on the apex of the follicle. These thick cells contain the rich rough endoplasmic reticulum, mitochondria, and Goldi bodies and possess many microvilli, indicating synthetic/secretory activities, and become thicker along with the oogenetic progress. Another region of epithelium does not exhibit these features. These results show the structural and functional modularity specific to some functions of the follicle epithelium of *H. japonica*. Therefore, the follicle epithelium of Myriapoda is divided into 3 types: physiologically functional uniformly, nonfunctional only, and those with both. We suggest the need to reconsider the nature and roles of ovarian somatic tissues of Myriapoda and Arthropoda.

## 1. INTRODUCTION

The ovarian somatic cells underpin female gametogenesis, i.e., oogenesis, in many metazoans (Matova and Cooley, 2001, Eckelbarger and Hodgson, 2021). The ovarian somatic tissue near oocytes, such as follicle epithelium, sometimes differentiates into some structural compartments related to the roles in oogenesis (e.g., Laale, 1980, Sugino et al., 1987, Büning, 1994, Morisawa, 1999, Kubrakiewicz et al., 2003, Plancha et al., 2005, Lambert, 2009). Especially many insect species have morphologically specialized follicle domains (Büning, 1994, Jaglarz et al., 2008, Tworzydlo and Kisiel, 2011). Margaritis et al. (1980) identified 11 cell subpopulations in the follicle epithelium of the fruit fly, *Drosophila melanogaster*. Each subpopulation has distinctive structural features in a specific follicle region and has a different role in oogenesis. For example, the posterior polar cells are round-shaped and persist in the posterior region of the follicle, which play roles in the differentiation of other follicle cells and the anteroposterior patterning of oocytes (Grammont and Irvine, 2002, Xi et al., 2003, Bastock and St Johnston, 2008, Roth and Lynch, 2009). The border cells are also round-shaped and initially placed on the anterior region of the follicle and then move with the anterior polar cells toward between oocytes and nurse cells, which contributes to the micropylar formation together with the anterior polar cells and the centripetal cells (Montell, 2003, Horne-Badovinac, 2020). The subpopulation referred to as the dorsolateral cells or the anterodorsal cells is initially a flat epithelial sheet but later becomes a tubular 3D structure and forms the dorsal appendage of the egg (Berg, 2005, Osterfield et al., 2013, Jouette et al., 2017). Hence, structural modularity is a strong indicator that the somatic cells are involved in oogenesis.

In Myriapoda, ovarian somatic cells around oocytes, i.e., the follicle cells, are a potential exception to the structural compartments and roles in oogenesis. The myriapod follicle epithelium is composed of simple thin cells. These cells are homogenous and show no synthetic/secretory activity (Crane and Cowden, 1968, Herbaut, 1974, Sareen and Adiyodi, 1983, Kubrakiewicz, 1991a, b, Miyachi and Yahata, 2012, Yahata et al., 2018). Also, the heterocellular junction between the follicle cells and the oocyte has not been found. Therefore, evidence of the structural and functional specialization of the ovarian somatic tissues was not obtained from myriapod species so far. Further, due to the absence of structural specialization and synthetic/secretory activity, the follicle cells are considered not to contribute to oogenesis in most myriapods investigated (briefly reviewed in Eckelbarger and Hodgson, 2021). The unique ovarian structure of Myriapoda would support this view: the basement membrane between the follicle epithelium and oocyte and oocyte growth within a hemocoelic space (Fig.1) (Kubrakiewicz, 1991a, Miyachi and Yahata, 2012, Yahata et al., 2018, Chikami and Yahata, 2019a), which is different from the oocyte growth within the ovarian lumen in other arthropods (Makioka, 1988, Büning, 1994, Kubrakiewicz et al., 2012, Jędrzejowska et al., 2014, Jędrzejowska, 2019). Thus, direct communication between the follicle cells and the oocyte may be prevented by the basement membrane.

**Fig. 1.**
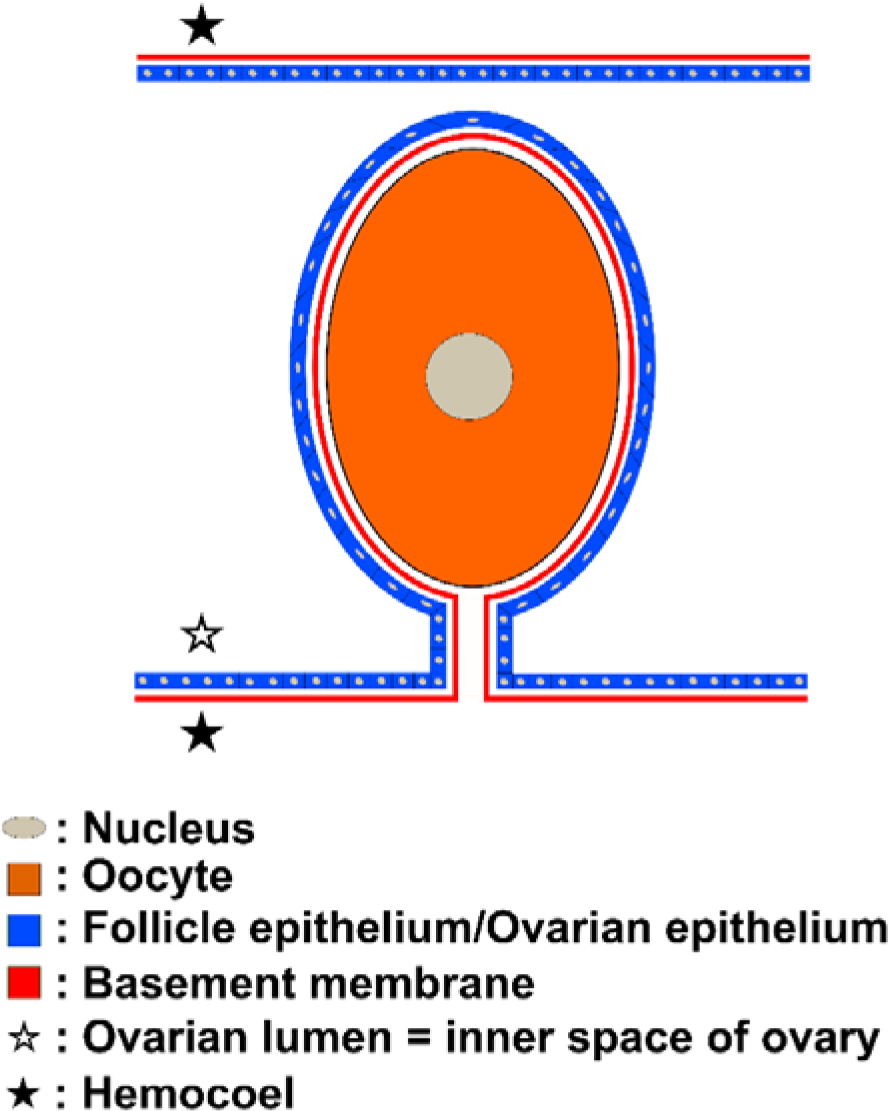
The ovarian structure of Myriapoda. The oocyte grows within the extraovarian environment, i.e., hemocoel. The basement membrane lines between the somatic cells and the oocyte.

In contrast to the conventional opinion on the structure and roles of the myriapod follicle epithelium, a recent study suggested potential structural and functional specialization of the follicle epithelium of Japanese pill millipedes, *Hyleoglomeris japonica* and *Hyleoglomeris yamashinai* (Chikami and Yahata, 2019b). The follicle epithelium is divided into two distinct regions by histological nature: the main-body region and the apical thick region. The former covers the majority of the follicle. The latter is consistently located on the apex of the follicle. These two regions are different in the height of the cells and sensitivities to phalloidin and eosin. These features suggest the presence of structural and functional heterogeneities in the follicle epithelium of myriapod species, facilitating the rethinking of the structure and function of the ovarian somatic tissues of Myriapoda. Understanding the ultrastructural and histochemical nature implicating some secretory activity and the correlation of the growth between these regions and oogenesis is required to verify the structural and functional modularity and to compare the insights between previous studies in other species and the follicle epithelium of *Hyleoglomeris* species.

In this study, we found differences in the ultrastructural nature within the follicle epithelium of *H. japonica*, indicating functional specialization. Also, the growth pattern was different between the regions of the follicle epithelium. Therefore, we reconsider the morphological nature of the follicle cells of Myriapoda and Arthropoda and propose a novel viewpoint and possible evolutionary history of the ovarian follicle.

## 2. Material and methods

### 2.1. Animals

135 individuals of *Hyleoglomeris japonica* Verhoeff, 1937 were collected in Mito City and Tsukuba City, Ibaraki Prefecture, and Minamisanriku Town, Miyagi Prefecture during May 2014–August 2016. The animals were kept in small plastic containers filled with leaf litter at 25°C.

### 2.2. Histological analysis

For histological analysis, dissected ovaries were fixed with Bouin’s solution (picric acid: formaldehyde: glacial acetic acid = 15: 5: 1) overnight or 4% paraformaldehyde (PFA) for 3 hours. For paraffin sectioning, the fixed specimens were dehydrated and cleared in a graded ethanol-n-butanol series, followed by embedded in paraffin (Sigma: Paraplast Embedding Media Paraplast Plus). The embedded specimens were cut into serial sections 5 µm thick using a microtome (American Optical: Model 820) with stainless knives (Feather: Microtome Blade Stainless Steel S35). For resin sectioning, the fixed specimens were dehydrated with an ethanol series before being embedded in the methacrylate resin (Kulzer: Technovit 7100). The embedded samples were cut into sections of 2 µm thickness. Both paraffin and resin sections were stained with Mayer’s hematoxylin and Eosin. Giemsa staining, azan staining, or alcian blue-periodic acid SCHIFF (PAS)-hematoxylin staining was also performed for the paraffin section. The stained sections were observed with light microscopies (Olympus: BH2 and Nicon: Optiphoto).

### 2.3. Ultrastructural analysis

For transmission electron microscopic analysis, the whole bodies were cut into some blocks (about 1.5 mm in length) and pre-fixed with 2.5% glutaraldehyde in 0.1 M phosphate buffer (PB) (pH 7.0–7.2) for 2h at room temperature. The pre-fixed specimens were washed with 0.1 M PB twice and post-fixed with 1% OsO4 in 0.1 M PB for 1 hour at 4°C, followed by washed with 0.1M PB twice for 1h. The post-fixed specimens were dehydrated in a graded acetone series and then embedded in Quetol 651 resin (Nisshin EM). The embedded specimens were cut into 60–80 nm thick ultrathin sections with an ultramicrotome (Leica: EM UC07). The sections were observed with transmission electron microscopy (Hitachi: H-7650).

### 2.4. Detection of nucleus and F-actin

For specifically staining the nucleus and F-actin, the ovarian tissues were fixed with 4% PFA for 3 hours. Then, the specimens were washed in phosphor buffered saline (PBS) (pH=7.0), followed by permeabilization with PBS with 0.5% Tween-20. The samples were stained by the 4’,6-diamidino-2-phenylindole (DAPI) and the rhodamine-conjugated phalloidin (ThermoFisher). The signals were detected by fluorescent microscopy (Nicon: Optiphoto).

### 2.5. Measurement and statistical analysis

To examine the correlation of the growth between the follicle cells and the oocyte, we measured the length of the major axis of the oocytes and the height of the main-body region and the apical thick region. First, we took photos of the ovarian follicles under differential interference contrast microscopy (Nicon: Optiphoto). Then, the oocyte length and the follicle regions’ height were measured from the photos using the Image J software (*N* = 51). We calculated the average of the three replications in each measurement. The quantitative data were listed in the supplementary material (Table S1). To evaluate the degree of the correlation of the growth between the follicle regions and the oocytes, we computed the Pearson product-moment correlation coefficient and conducted the correlation test using the *cor.test* function of the R-v4.2.3 (R Core Team, 2023). We took *P*-value < 0.05 as the significant level.

## 3. RESULTS

### 3.1. Basic structure of the ovary and growth position of oocytes of *H. japonica*

The ovarian structure of *H. japonica* was observed at the light microscopic level (Yahata and Makioka, 1997, Chikami and Yahata, 2019b), which suggests that its oocytes grow outside the ovary. However, the position of the basement membrane, which is essential for verifying the oocyte growth position, has yet to be investigated. First, we confirm this assumption at the ultrastructure level.

The ovarian follicle of *H. japonica* comprised single-layered epithelial cells (Fig. 2A, B). This epithelial layer continued to the ovarian epithelium constituting the ovarian wall (Fig. 2B, C). Both ovarian epithelium and follicle epithelium was lined with its basement membrane (Fig. 2D, E). Hemidesmosomes could be detected between the cell and its basement membrane (Fig. 2E). Also, the basement membrane of the follicle epithelium continued to that of the ovarian epithelium (Fig. 2D). Thus, the follicle of *H. japonica* was regarded as a pocket opened into the hemocoel, consisting with the histological observation by Yahata and Makioka (1997). Then, the basement membrane separates oocytes from the follicle cells. There were hemocytes and tracheoles in the space between the oocyte and the basement membrane (Fig. 2D, F, G). Therefore, the oocytes of *H. japonica* grew within the hemocoelic environment, which strongly supports the hypothesis that the hemocoelic nature of oocyte growth is a synapomorphy in Myriapoda (Yahata et al. 2018).

**Fig. 2.**
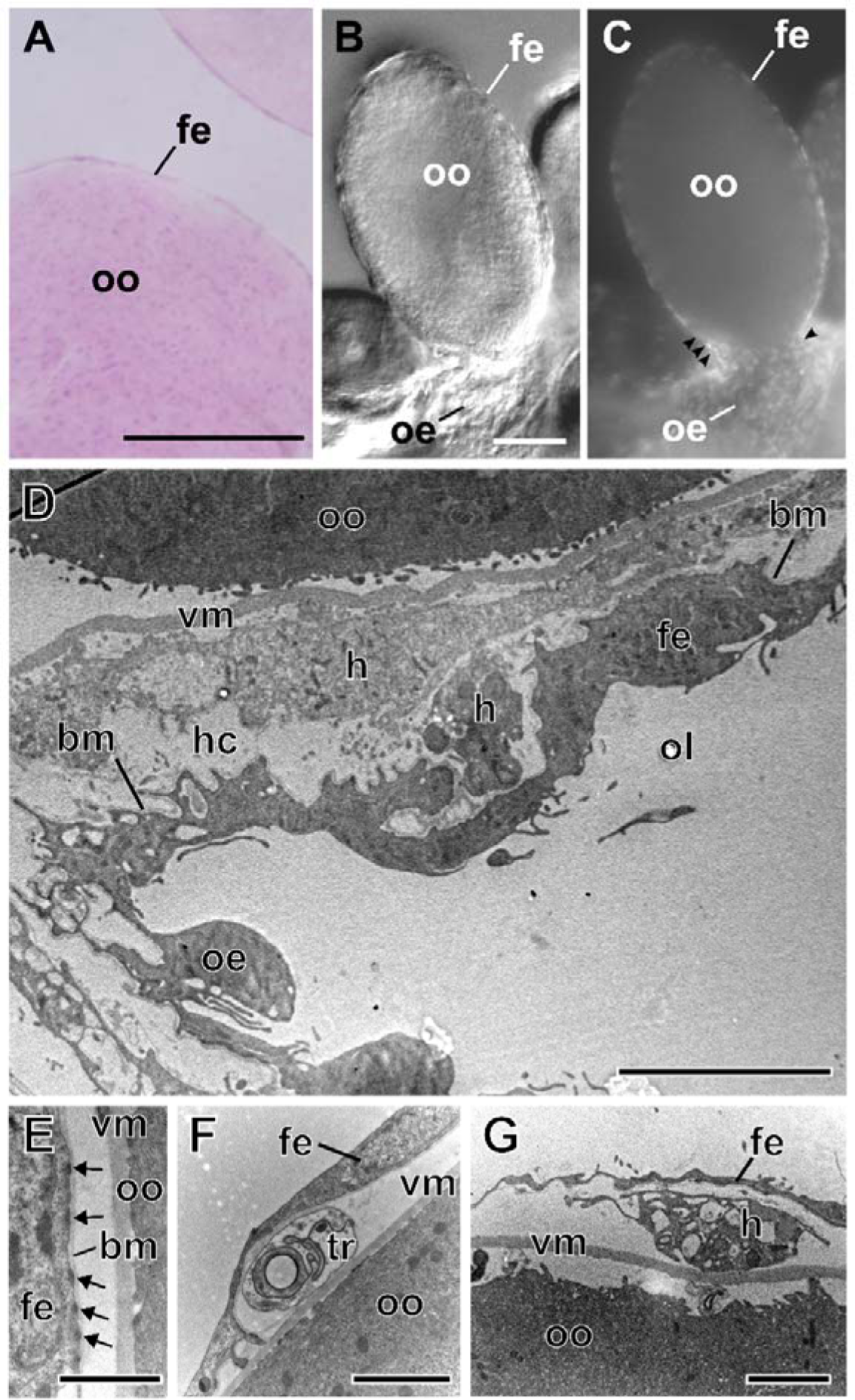
The basic structures of the ovary in *H. japonica*. (A–C) the ovarian follicle at the light microscopic level. (A) Resin section with HE staining. (B) DIC image. (C) Nucleus detection by DAPI staining. The arrowheads indicate the contact zone between the ovarian epithelium and the follicle epithelium. (D) The contact zone between the ovarian epithelium and the follicle epithelium. (E) The enlargement view of the basal side of the follicle cell. Arrows indicate hemidesmosomes. (F, G) The tracheal cell (F) and hemocyte (G) between the follicle epithelium and oocyte. Scales: 2 µm. bm, basement membrane; fe, follicle epithelium; h, hemocyte; hc, hemocoel; oe, ovarian epithelium; ol, ovarian lumen; oo, oocyte; tr, tracheal cell; vm, vitelline membrane. Scales: 50 µm (A, B), 5 µm (D), 1 µm (E), 2 µm (F, G).

### 3.2. Ultrastructural nature of the compartments of follicle epithelium of *H. japonica*

To investigate the structural differences between the two regions in the follicle epithelium (Fig. 3A) of *H. japonica* mentioned by Chikami and Yahata (2019b), we observed the ultrastructure of these regions.

**Fig. 3.**
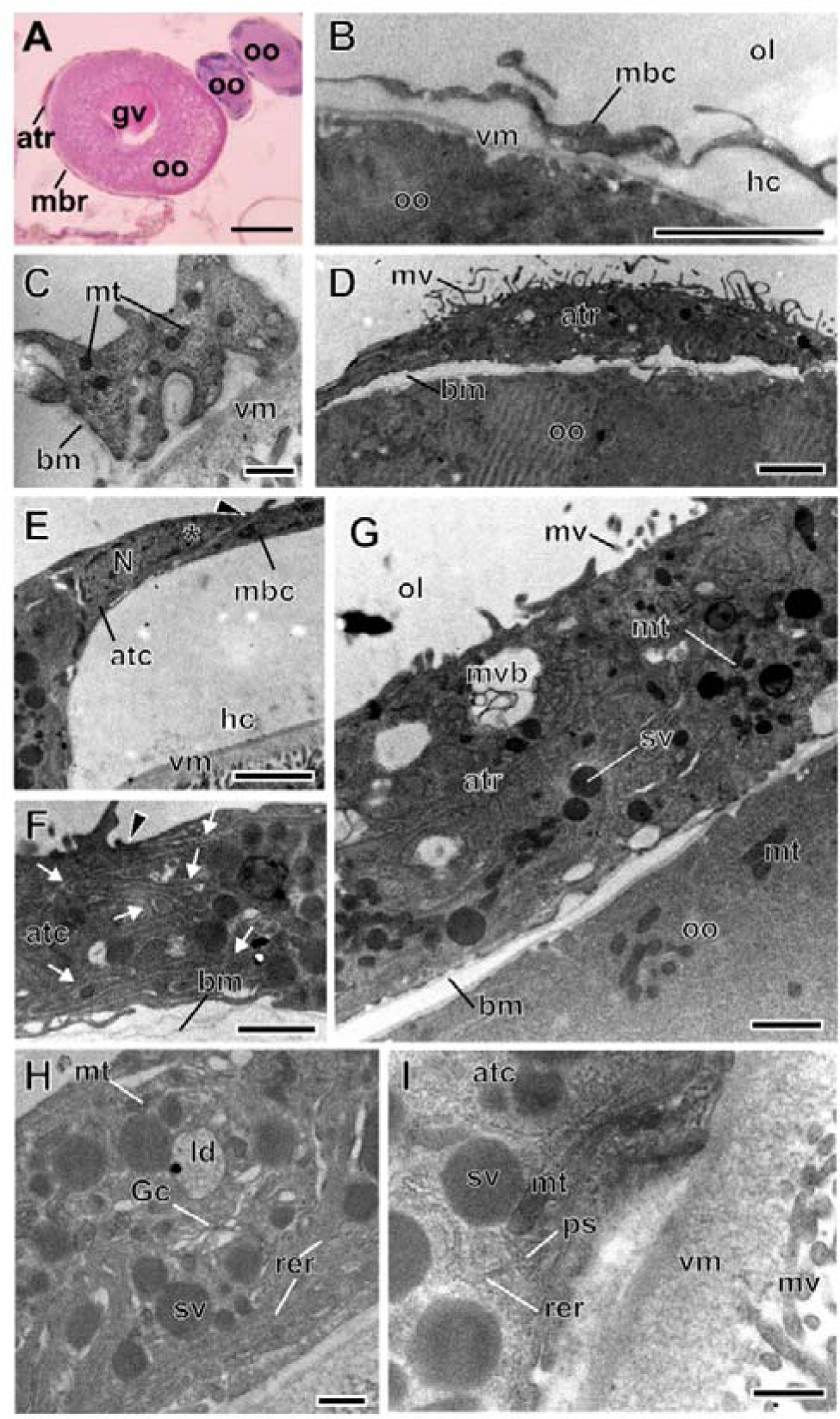
The structural nature of the two regions of the follicle epithelium in *H. japonica*. (A) The gross view of the ovarian follicles. Paraffin section. HE staininig. (B, C) The low (B) and high (C) magnified images the main-body epithelium. (D) The low magnified view of the apical thick region. (E) The contact between the main-body region cell and the apical thick region cell. The arrowheads and asterisk indicate the adherence and septate junction, respectively. (F) The adhesion zone between cells of the apical thick region. The arrows indicate the intercellular space. (G–I) The organelles within the apical thick region. The low (G), middle (H), and high (I) magnified images of the apical thick region cell. atc, apical thick region cell; atr, apical thick region; bm, basement membrane; Gc, Goldi complex; gv, germinal vesicle; hc, hemocoel; ld, lipid droplet; mbc, main-body region cell; mbr, main-body region; mt, mitochondria; mv, microvilli; mvb, multivesicular body; N, nucleus; ol, ovarian lumen; oo, oocyte; ps, polysome-like structure; rer, rough endoplasmic reticulum; sv, secretory vesicle; vm, vitelline membrane. Scales: 50 µm (A, D, E), 2 µm (B), 1 µm (G), 0.5 µm (C, F, H, I).

The main-body region was constituted of single-layered flattened epithelial cells (Fig. 3B). These cells were systematically lined by the basement membrane (Fig. 3C). Also, the apical side of the epithelium had no or a few microvilli. The main-body cell had some round-shaped mitochondria (Fig. 3C). However, we could detect no or few active organelles such as the Goldi body, endoplasmic reticulum, and endo/exocytotic vesicles (Fig.3B, C). The cytoplasm of the main-body cell was thus homogenous and showed a low synthetic/secretory activity.

The apical thick region was a columnar epithelial layer (Fig. 3D). This region continued to the main-body region with the adherence and septate junctions (Fig. 3E). The cells of this region also possessed the basement membrane on their basal side (Fig. 3D, F). The epithelium was not a single simple layer but a complicatedly folded pseudostratified one (Fig. 3D, F). The microvilli were enriched on the apical surface of the cells (Fig. 3F). Further, there were a significant number of secretory vesicles, rough endoplasmic reticulum, polysome-like structures, elongated mitochondria, Goldi complexes, and lipid droplets in the cytoplasm of the apical thick region (Fig. 3G, H, I). Also, we detected some multivesicular bodies (Fig. 3G), which suggests some intracellular digestion. Thus, the cytoplasm of the apical thick region had some metabolic activities.

### 3.3. Histochemical nature of the apical thick region

The ultrastructure of the follicle epithelium of *H. japonica* showed rich synthetic/secretory activities in the apical thick region. Chikami and Yahata (2019b) consistently show that the apical thick region is more intensely stained with the eosin than the main-body region. Thus, the apical thick region could contain rich proteins such as secretory proteins. Here, we investigated the histochemical features to verify this possibility.

First, we confirmed the eosinophilicity of the apical thick region and revealed that the cytoplasm of the region was strongly stained by eosin (Fig. 4A). Then, we performed azan staining to diagnose types of secretory granules. The cytoplasm of the apical thick region was red-colored (Fig. 4B). Also, some small granular structures were stained in blue color (Fig. 4B). To investigate granular proteins, we used the PAS staining. The apical thick region showed stronger PAS-positive reactivity, which suggests the secretory proteins (Fig. 4C). We could not detect these features in the main-body region. To examine the richness of nucleic acids, we observed the apical thick region stained by hematoxylin or Giemsa’s solution. It was hard to detect hematoxylin-positive signals (Fig. 4A). However, some purplish-red subregions in the apical thick region were stained by Giemsa’s solution, suggesting the rich nucleic acids (Fig. 4D).

**Fig. 4.**
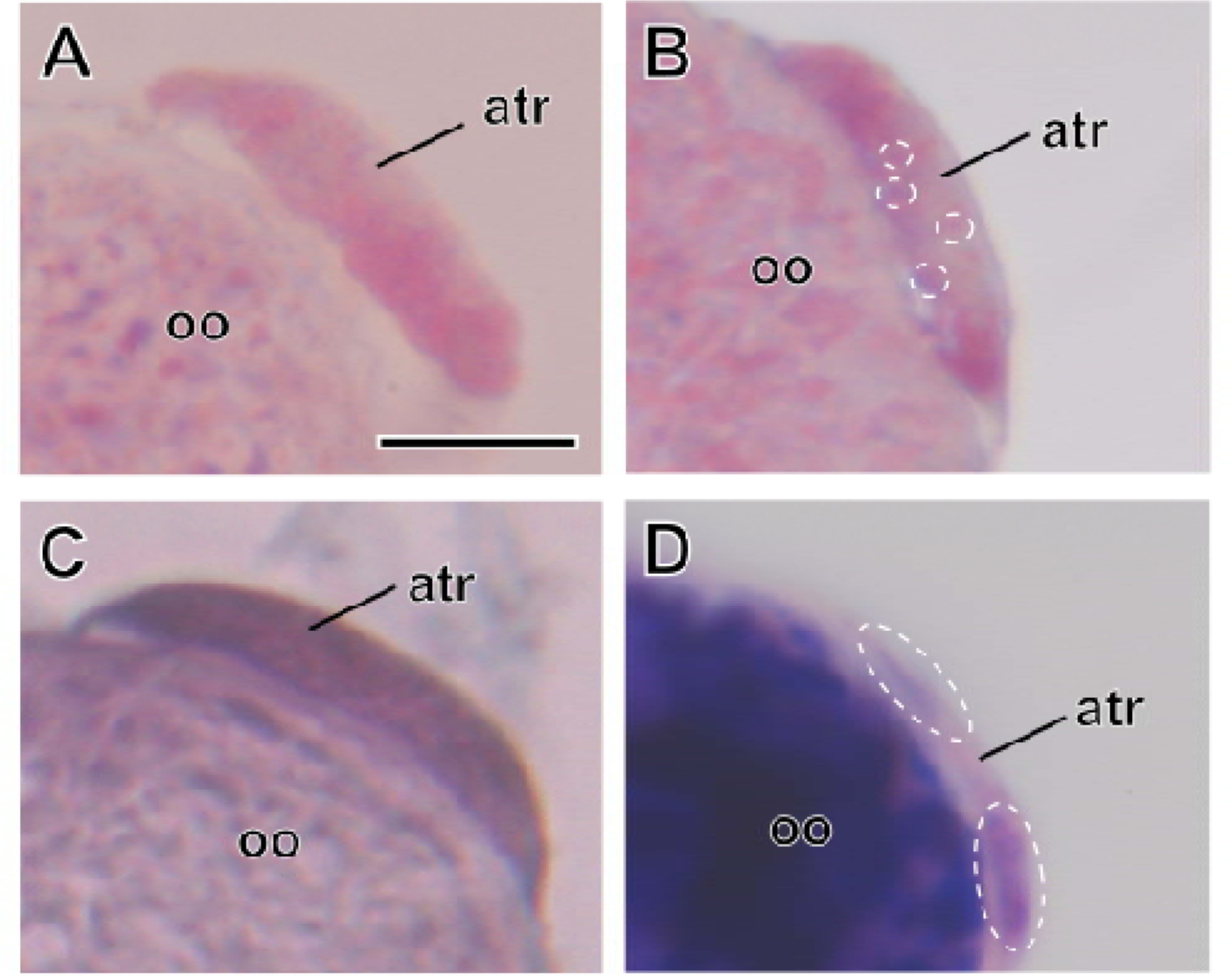
The histochemical nature of the apical thick region in the previtellogenesis. (A) HE staining. (B) Azan staining. (C) Alcian blue-PAS-hematoxylin staining. (D) Giemsa staining. The dotted lines indicate the secretory proteins (B) and the nucleic acids (D), respectively. Paraffin section. atr, apical thick region; oo, oocyte. Scale: 10 µm.

### 3.4. Development of the follicle epithelial cells during the oogenesis

If the region contributes to some aspects of oogenesis, it is assumed that the growth of the apical thick region is associates with the oogenesis. To investigate this prediction, we observed the development of the apical thick region during the oogenesis.

First, we examined when the apical thick region is morphologically differentiated from another region. We could not find any morphologically distinguished region in the apex of the early previtellogenic follicle when the oocytes grew to 50 µm in length and were round (Fig. 5A). During the middle previtellogenesis, when the oocytes were 50–120 µm and showed an oval shape, about 40% of the follicle epithelium remained undifferentiated (Fig. 5B). In contrast, the rests already had a thicker domain on the apex of the follicle (Fig. 5C). In the late previtellogenesis, when the oocytes grew above 125 µm, the morphologically distinguishable structure could be found on the apexes of all follicles (Fig. 5D). The morphologically detectable region persistently showed phalloidin susceptibility (Fig. 5C’, D’) as described in Chikami and Yahata (2019b). Thus, the uniformity of the follicle epithelium was first morphologically broken during the middle previtellogenesis.

**Fig. 5.**
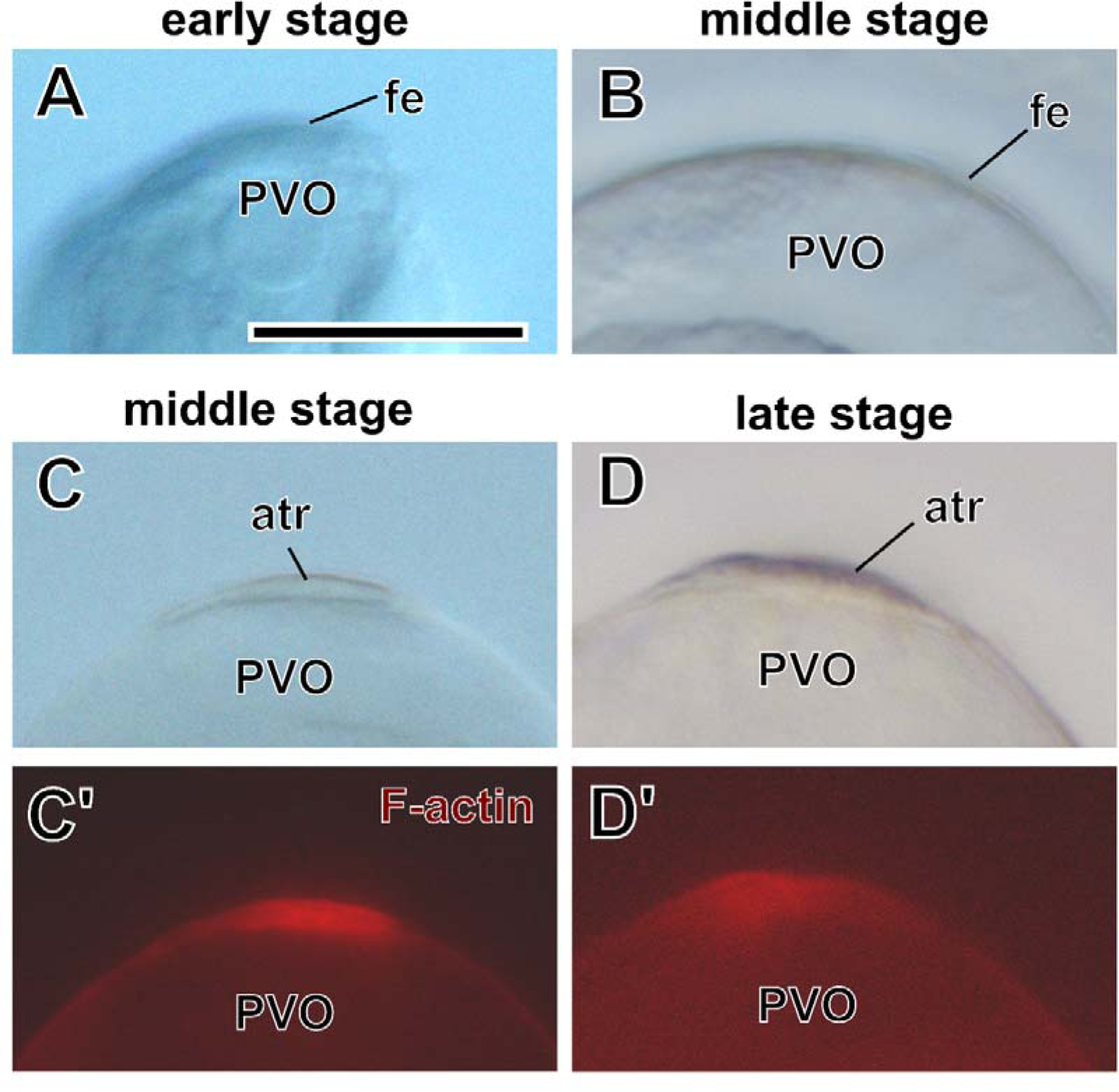
The differentiation into the apical thick region. (A) During the early previtellogenesis, no morphologically visible structure can be found. In middle previtellogenesis, the apex of the follicle epithelium of some follicles remains the thin layer (B) but other follicles have the morphologically visible thick region (C) which shows phalloidin subspeciality (C’). Finally, the apical thick region can be detected morphologically (D) and phalloidin-positively (D’) in all follicles during the late vitellogenesis. atr, apical thick region; fe, follicle epithelium; PVO, previtellogenic oocyte. Scale: 30 µm.

Second, we investigated growth correlations between the two follicle compartments and the oocytes. If a follicle epithelial region is involved in the oogenesis, the region may become thicker along with the oogenesis. Then, we revealed that the apical thick region gradually thickened during the previtellogenesis (Pearson product-moment correlation coefficient *r* = 0.78, *t* = 8.72, df = 49, *P* = 1.53×10^-11^, Fig. 6). We could not detect a significant effect on the correlation between the main-body region and the oocyte growth (Pearson product-moment correlation coefficient *r* = 0.20, *t* = 1.44, df = 49, *P* = 0.16, Fig. 6). Also, the correlation coefficient *r* of the main-body region was lower than that of the apical thick region. Thus, the main-body region did not grow much during the previtellogenesis. In contrast, the apical thick region significantly grew along the previtellogenesis.

**Fig. 6.**
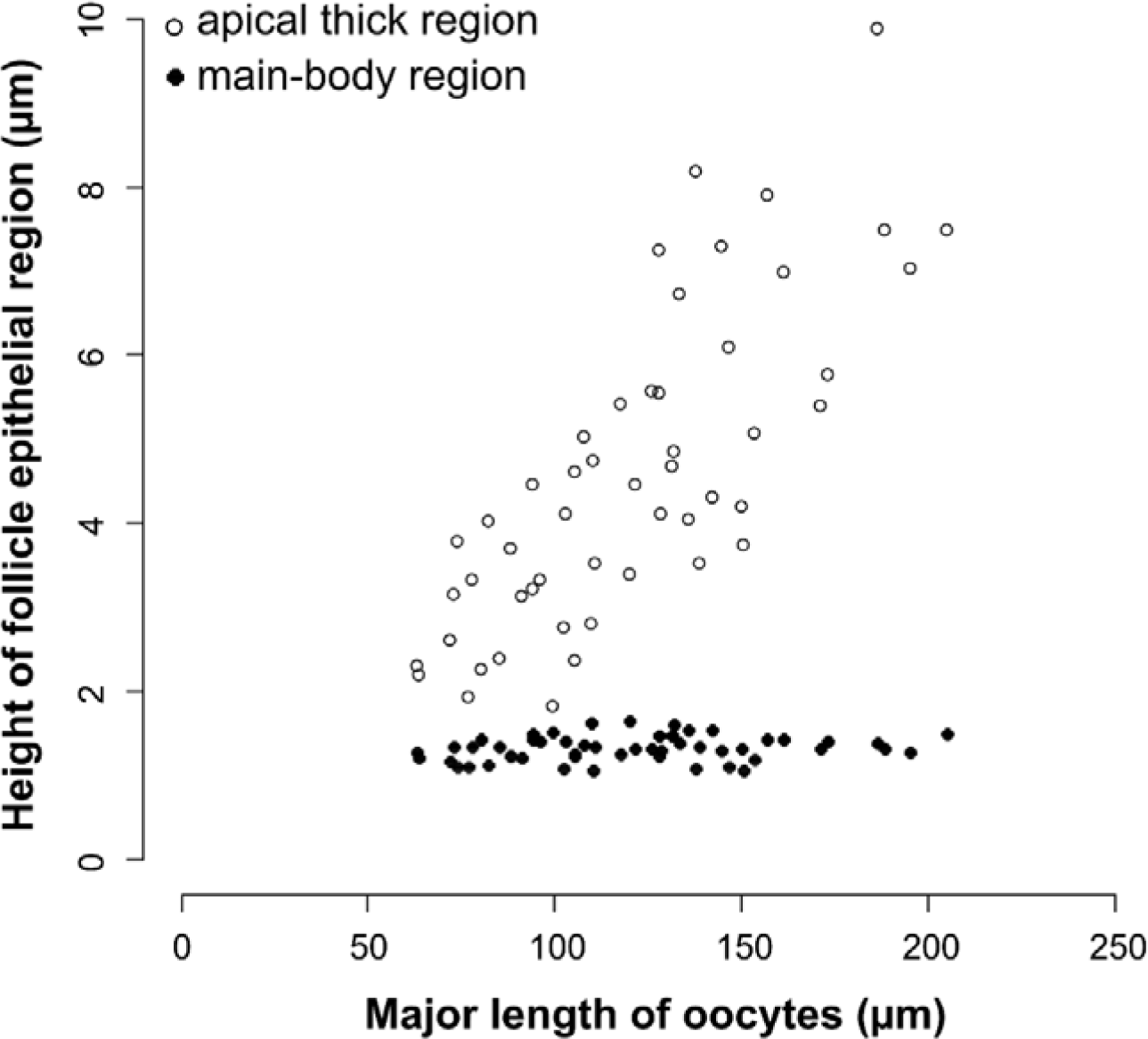
The correlation of growth between the follicle epithelial regions and the oocytes. The Pearson product-moment correlation coefficient *r* is 0.78 and 0.20 in the apical thick region and the main-body region, respectively. *N* = 51. The data used for this figure is described in Table S1.

Finally, we observed the morphological nature of the apical thick region during the vitellogenesis. The apical thick region persisted during the vitellogenesis and was likely to become stretched (Fig. 7A, B). The organelles of the apical thick region during the vitellogenesis were similar to those during the previtellogenesis. However, the basement membrane in the vitellogenesis was partially disrupted and became hard to be visible (Fig. 7C) in contrast to that in previtellogenesis (Fig. 7D). Consistently, the hemidesmosomes between the epithelium and its basement membrane became sparse in the disrupted area. Also, the electron-dense substance accumulated between the follicle cells and the oocyte (Fig. 7C).

**Fig. 7.**
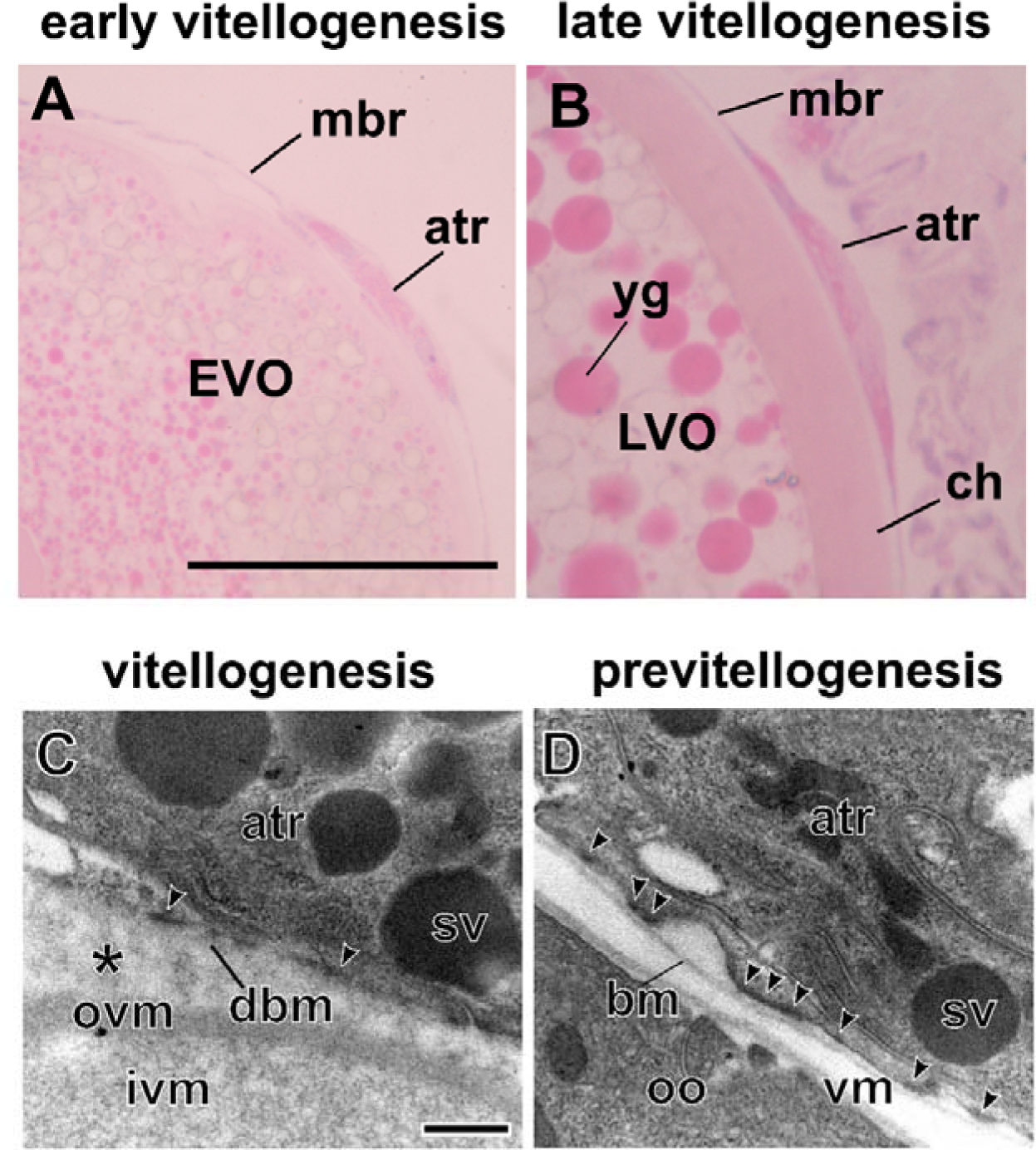
Morphological nature of the apical thick region during oogenesis. (A, B) The light microscopic images of the apex of the follicle during early (A) and late (B) vitellogenesis. Resin section. HE staining. (C, D) The ultrastructure of the basal side of the apical thick region cell in vitellogenesis (C) and previtellogenesis (D). The arrowheads indicate the hemidesmosomes between the cell and its basement membrane. The asterisk shows the electron-dens substance. atr, apical thick region; bm, basement membrane; ch, chorion; dbm, degenerative basement membrane; EVO, early vitellogenic oocyte; ivm, inner vitelline membrane; LVO, late vitellogenic oocyte; Mbr, main-body region; oo, oocyte; ovm, outer vitelline membrane; sv, secretory vesicle; vm, vitelline membrane; yg, yolk granule. Scales: 50 µm (A), 0.5 µm (C).

## 1.4. DISCUSSION

### 4.1. Structural and functional modularity of the follicle epithelium of *H. japonica* and Arthropoda

Prior to this study, two distinct regions were identified in the ovarian follicle epithelium in two *Hyleoglomeris* species based on the differences in the histological nature at the light microscopic level (Chikami and Yahata, 2019b). Additionally, this study discovered the differences in the ultrastructural nature and developmental manner between these regions (Fig. 8). Our findings show that the apical thick region is not just a swelling cell mass but is a morphological/functional unit. Therefore, the main-body and apical thick regions can be regarded as structural and functional modularity in the ovarian follicle epithelium in *H. japonica*.

**Fig. 8.**
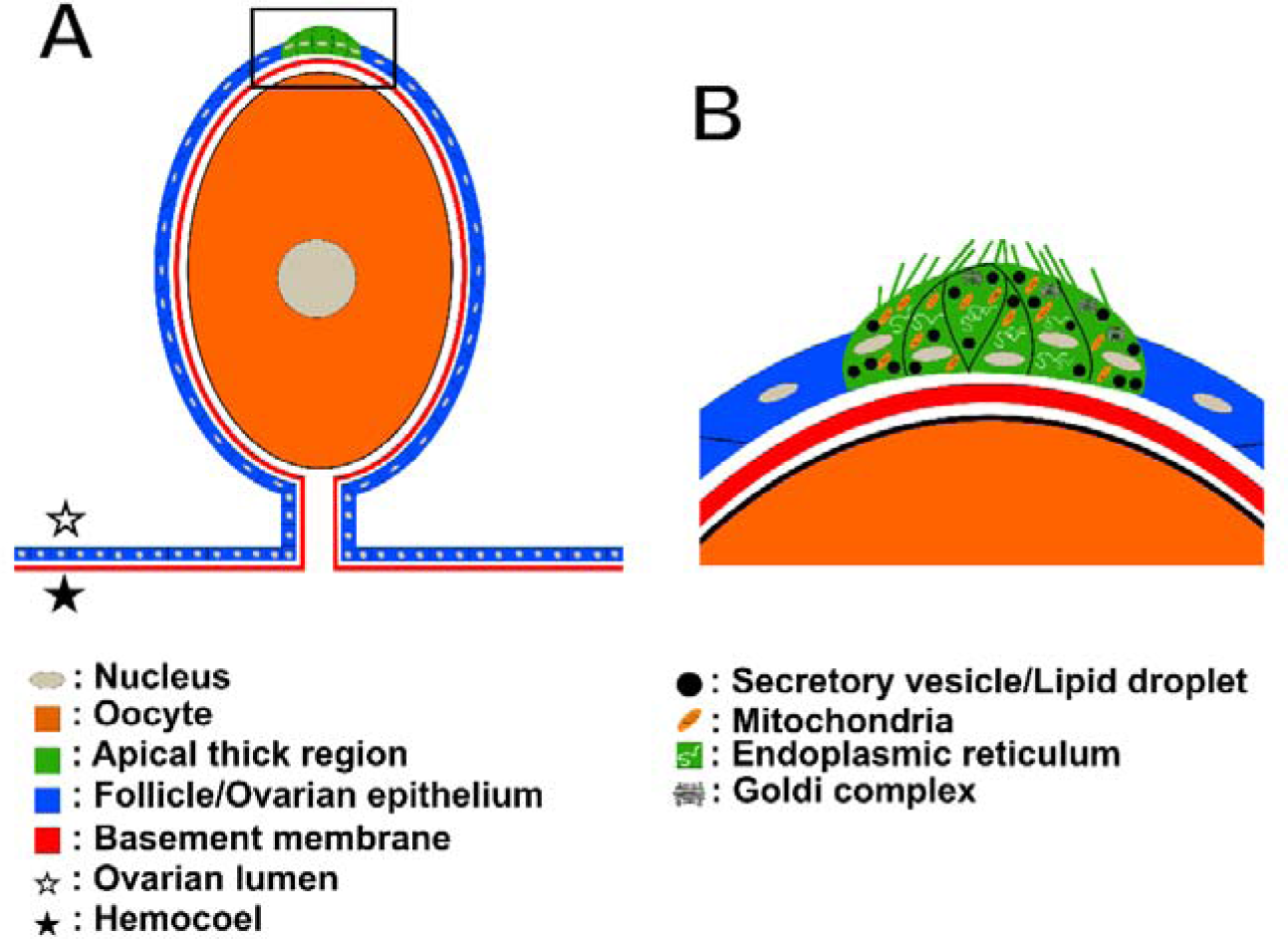
The schematic drawing of the structural modularity of the follicle epithelium in *H. japonica*. (A) The overall follicle of *H. japonica*. (B) The enlargement view of the apical thick region indicated by the frame in (A).

The structural and functional modularity within a somatic tissue covering the oocyte has been known from the follicle cells of many insects such as Diptera (Büning, 1994, Mazurkiewicz and Kubrakiewicz, 2005), Lepidoptera (Yamauchi and Yoshitake, 1984a, b, Mazurkiewicz-Kania et al., 2019), Neuroptera (Kubrakiewicz et al., 2005, Garbiec et al., 2016), Psocodea (Zawadzka et al., 1997), Hemiptera (Büning, 1994), Dermaptera (Tworzydlo and Kisiel, 2011), and Mantophasmatodea (Tsutsumi et al., 2005). Over a century, many histological and ultrastructural studies have uncovered the morphological features of myriapod ovaries and oogenesis (e.g., Tönniges, 1902, Tiegs, 1940, 1945, 1947a, b, Jangi, 1957, Herbaut, 1974, Knoll, 1974, Biliński, 1979, Sareen and Adiyodi, 1983, Kubrakiewicz, 1991a, b, Yahata and Makioka, 1994, 1997, Fontanetti et al., 2010, Miyachi and Yahata, 2012, Yahata, 2012, Yahata et al., 2018, Chikami and Yahata, 2019a). However, since the follicle epithelium has been described as a simple and homogenous tissue without active organelles, the structural and functional specialization of the follicle epithelium has not been reported from Myriapoda. Also, to our knowledge, the evident structural and functional modularity of a single somatic tissue surrounding the oocyte has not been reported from Chelicerata and crustaceans (e.g., Jaglarz et al., 2020). Thus, the main-body and apical thick region of *H. japonica* would provide a novel viewpoint on the structural and functional specialization of the follicle epithelium in arthropods other than insects and Myriapoda.

### 4.2. Functions of the follicle epithelium of *H. japonica*

The main-body region of the follicle epithelium in *H. japonica* lacked metabolic activities. The role of the ovarian somatic tissues without secretory activities is the mechanical support of follicles (Eckelbarger and Hodgson, 2021). Therefore, it is plausible that the main-body region of *H. japonica* bears the support of the follicle structure as same as the follicle cells in most myriapods investigated (Kubrakiewicz, 1991a, b).

The apical thick region contained many active organelles and microvilli and showed distinctive histochemical nature indicating protein synthesis and modification. In this study, specific roles of the apical thick region still need to be clarified as the experimental verification is prevented by the current technical limitation in the ablation and the ex vivo or in vitro culture of the follicle cells. The growth of the apical thick region following the progression of the oogenesis and the degeneration of its basement membrane during vitellogenesis however suggests the possibility that the follicle cells of *H. japonica* contribute to some physiological roles in the oogenesis. In many insects, the specialized follicle subpopulations are involved in the formation of special structures of the egg, such as the micropyle and dorsal appendage (Büning, 1994, Suzanne et al., 2001). However, no evidence has been obtained of such special structures in *Hyleoglomeris* and other myriapod eggs. Since the apical thick region features would be partially similar to the follicle cells of the second generation in the proturan, *Acerentomon gallicum*, which are involved in choriogenesis (Bilinski and Klag, 1977, Bilinski, 1994), chorion formation is a candidate for the role of the apical thick region. A future task will be to reveal the relationship between the specialized follicle epithelium and the oogenesis of *H. japonica*.

### 4.3. Reconsideration of the morphological plan of the somatic tissue around the oocyte of Myriapoda

Most ultrastructural studies on myriapod oogenesis suggest that the myriapod oogenesis is a solitary manner due to the absence of physiological activities of the follicle cells (e.g., Herbaut, 1974, Sareen and Adiyodi, 1983, Kubrakiewicz, 1991a, Miyachi and Yahata, 2012, Yahata et al., 2018). Kubrakiewicz (1991a, b) called such follicle epithelium in a julid millipede, *Ophyiulus pilosus,* an ‘ovarian sac epithelium’ and distinguished it from that of the Hexapoda because this epithelium did not have any synthetic/secretory activity and any structural specialization. This view is consistent with the hemocoelic nature of the oocyte growth in Myriapoda (Kubrakiewicz, 1991a, Miyachi and Yahata, 2012, Yahata et al., 2018), which allows the oocyte to obtain nutrients from the blood directly. Also, this notion does not contradict the synthesis of yolk precursors in hemolymph and the fat body in Myriapoda (Prasath and Subramoniam, 1991). Exceptionally, although the structural and functional specialization has not been described, the ultrastructural study suggests that the ovarian somatic cells surrounding the oocyte of the bristle millipede, *Polyxenus lagurus*, are involved in its vitellogenesis (Kubrakiewicz, 1991c), which can be referred to as the ‘true’ follicle epithelium in terms of the function. Like *P. lagurus*, it is reasonable that the ovarian somatic tissue around the oocyte of *H. japonica* is the true follicle epithelium. The conventional view and our findings indicate at least three types of somatic epithelium around oocytes in Myriapoda: the uniform true follicle epithelium with physiological function, the uniform ovarian sac epithelium without the function, and the epithelium bearing both structurally divided functional and nonfunctional regions.

*P. lagurus* belongs to Polyxenida, the most basal clade of Diplopoda (Blanke and Wesener, 2014, Miyazawa et al., 2014, Rehm et al., 2014, Fernández et al., 2016, 2018, Rodriguez et al., 2018, Benavides et al., 2023). Also, *H. japonica* investigated in this study is a species of Pentazonia, the sister group of other Chilognatha (= diplopods other than Polyxenida). In contrast, species with the ‘follicle epithelium’ that seems not to participate in oogenesis, such as *O. pilosus*, are usually classified into more derived diplopod taxa. Therefore, this phylogenetical distribution implies that the physiologically functional follicle epithelium may be a plesiomorphic feature of Diplopoda and that the roles in the oogenesis were secondarily lost during diplopod evolution. We speculate that the structural specialization of *H. japonica* might represent an intermediate state during the gradual evolution from the uniform functional mode in the true follicle epithelium to the completely nonfunctional epithelium, i.e., the ovarian sac epithelium. Alternatively, the structural/functional specialization appeared during the evolution from the common ancestor of Pentazonia to *Hyleoglomeris*. It might have facilitated some specific requirements for the oogenesis of *Hyleoglomeris* species, such as the primary axial formation.

Overall, we provided the first example of the structural/functional specialization of the myriapod follicle epithelium and proposed its potential evolutionary routes. Our results will facilitate reconsidering the morphological plan of the myriapod follicle epithelium. Like many myriapods, the ovarian somatic tissues surrounding oocytes in some other arthropods are suggested to be morphologically uniform (e.g., Jaglarz et al., 2020) or insignificant for the oogenesis (e.g., Jędrzejowska et al., 2013). The morphological studies on follicle epithelium’s structural/functional specialization in broad arthropod taxa will better understand the evolution and ground plan of the follicle epithelium in Myriapoda and Arthropoda.

## CRediT author statement

**Yasuhiko Chikami:** Conceptualization, Methodology, Formal analysis, Investigation, Resources, Writing – Original Draft, Visualization. **Kensuke Yahata:** Conceptualization, Methodology, Resources, Writing – Review & Editing.

## Declarations of interest

The authors declare that there is no conflict of interest in this study.

## Funding

This research did not receive any specific grant from funding agencies in the public, commercial, or not-for-profit sectors.

## Acknowledgements

We express our thanks to Dr. H. Mitsumoto, Mr. T. Suguro, Ms. E. Umetani, Ms. M. Shibata, Ms. Y. Takatani, Mr. N. Naya for encouragement to collect *Hyleoglomeris japonica*. Our analysis of transmission electron microscope was carried out with instruments at Open Facility, Research Facility Center for Science and Technology, University of Tsukuba.

## Data availability

All data used for statistical analysis is listed in the supplementary material. Other data will be made available on request.

## Notes

### Competing Interest Statement

The authors have declared no competing interest.

